# Cryptic splicing in synaptic and membrane excitability genes links TDP-43 loss to neuronal dysfunction

**DOI:** 10.1101/2025.08.28.672801

**Authors:** Caiwei Guo, Kuchuan Chen, Sarat C. Vatsavayai, Tetsuya Akiyama, Yi Zeng, Chang Liu, Odilia Sianto, Edith Yang, Juliane Bombosch, Rasheen Powell, Shannon Zhen, Shila Mekhoubad, Ryan D. Morrie, Georgiana Miller, Eric M. Green, Leonard Petrucelli, William W. Seeley, Aaron D. Gitler

## Abstract

TDP-43 pathology is a defining pathological hallmark of multiple neurodegenerative diseases, including amyotrophic lateral sclerosis (ALS) and frontotemporal dementia (FTD). A major feature of TDP-43 pathology is its nuclear depletion, leading to the aberrant inclusion of cryptic exons during RNA splicing. *STMN2* and *UNC13A* have emerged as prominent TDP-43 splicing targets, but the broader impact of TDP-43-dependent cryptic splicing on neuronal function remains unclear. Here, we report new TDP-43 splicing targets critical for membrane excitability and synaptic function, including *KALRN, RAP1GAP, SYT7* and *KCNQ2*. Using human stem cell-derived neurons, we show that TDP-43 reduction induces cryptic splicing and downregulation of these genes, resulting in impaired excitability and synaptic transmission. In postmortem brains from patients with FTD, these cryptic splicing events occur selectively in neurons with TDP-43 pathology. Importantly, suppressing individual cryptic splicing events using antisense oligonucleotides partially restores neuronal function, and combined targeting almost fully rescues the synaptic deficit caused by TDP-43 loss. Together, our findings provide evidence that cryptic splicing in these synaptic and membrane excitability genes is not only a downstream marker but instead a direct driver of neuronal dysfunction, establishing a mechanistic link between TDP-43 pathology and neurodegeneration in ALS and FTD.

## Introduction

Amyotrophic lateral sclerosis (ALS) and frontotemporal dementia (FTD) are two neurodegenerative diseases that affect different areas of the nervous system but share significant overlap in clinical, pathological, and mechanistic features (Ling et al., 2013). A key shared hallmark is the depletion of the RNA-binding protein TDP-43 from the nucleus and the formation of cytoplasmic inclusions, collectively referred to as TDP-43 pathology, which is observed in ∼97% of patients with ALS and ∼45% of patients with FTD (Arai et al., 2006; Neumann et al., 2006). In normal cells, TDP-43 is predominantly localized in the nucleus, where it is essential for various RNA metabolic activities, including transcription, pre-mRNA splicing, RNA stability, and RNA transport (Lagier-Tourenne et al., 2010).

Pre-mRNA splicing is a crucial step in gene expression, particularly in the brain, which has the highest level of alternative splicing among all tissues, with over 40% of neuronal genes undergoing at least one alternative splicing events (Yeo et al., 2004). A specific type of splicing involves the inclusion of non-canonical, or cryptic, exons and TDP-43 has emerged as a critical repressor of these cryptic RNA splicing events (Ling et al., 2015). When TDP-43 is depleted from the nucleus, cryptic exons that are normally excluded are incorporated into mRNAs, which can lead to RNA degradation or translation errors due to an open reading frame shift, resulting in diminished or non-functional protein products. Additionally, some cryptic splicing transcripts may generate *de novo* cryptic peptides that accumulate in the cerebrospinal fluid of ALS/FTD patients, offering potential as diagnostic biomarkers (Calliari et al., 2024; Irwin et al., 2024; Seddighi et al., 2024).

TDP-43’s function as a cryptic splicing repressor is conserved, but its specific targets are not (Ling et al., 2015). One TDP-43 cryptic splicing target in humans is *Stathmin-2* (*STMN2*), a gene necessary for axonal growth and regeneration (Klim et al., 2019; Melamed et al., 2019). Loss of TDP-43 promotes the inclusion of a cryptic exon in *STMN2*, which results in premature polyadenylation and a truncated non-functional mRNA, reducing its protein expression and impairing axonal regeneration. Importantly, suppressing the cryptic splicing event in human induced pluripotent stem cell (iPSC)-derived motor neurons can restore *STMN2* expression and axonal regeneration, suggesting that *STMN2* cryptic splicing contributes to enhanced neuronal vulnerability in the absence of TDP-43. Based on these findings, *STMN2* antisense oligonucleotides (ASOs) have been developed to block *STMN2* cryptic splicing and restore its protein level despite TDP-43 loss in humanized *STMN2* mouse models (Baughn et al., 2023).

Another TDP-43 cryptic splicing target is *UNC13A*, a major risk gene for ALS/FTD (Brown et al., 2022; Ma et al., 2022). Loss of TDP-43 from the nucleus causes a cryptic exon to be included in the *UNC13A* mRNA, resulting in nonsense-mediated mRNA decay and loss of UNC13A protein. Genetic variants associated with ALS/FTD in humans are located nearby the cryptic exon and promote cryptic exon inclusion when TDP-43 is dysfunctional (Brown et al., 2022; Ma et al., 2022; van Es et al., 2009). UNC13A plays a critical role in the nervous system in regulating synaptic vesicle fusion (Augustin et al., 1999; Böhme et al., 2016). Loss of TDP-43 in human neurons causes defects in synaptic transmission and neuronal activity, which, importantly, can be rescued by suppressing *UNC13A* cryptic exon inclusion (Keuss et al., 2024).

Beyond *UNC13A* and *STMN2*, do other TDP-43 cryptic splicing targets contribute to FTD/ALS pathogenesis and represent new potential biomarkers or therapeutic targets? In our previous study in which we identified *UNC13A* as a TDP-43 splicing target, we reported 66 genes that exhibit cryptic RNA splicing in the human brain owing to nuclear TDP-43 depletion (Ma et al., 2022). Many of these cryptic exons are in genes that are critical for neuronal function and therefore may contribute to neurodegeneration. Here we expand our analysis and demonstrate that cryptic splicing events in multiple membrane excitability and synaptic genes are direct consequences of TDP-43 loss, leading to decreased gene expression. Furthermore, we observe these cryptic splicing events in TDP-43-depleted neurons in the postmortem frontal cortex of patients with FTD. By using multielectrode arrays (MEAs) to monitor neuronal activity, we show that these genes encode proteins essential for membrane excitability and synaptic function. Inhibiting individual cryptic splicing events with ASOs partially restores the synaptic dysfunction resulting from TDP-43 reduction and simultaneously blocking multiple cryptic splicing events largely rescues it. Taken together, these findings provide evidence that cryptic splicing in multiple synaptic and membrane excitability genes is a key mechanism underlying TDP-43–associated pathology and expand the portfolio of TDP-43-regulated cryptic splicing targets in FTD/ALS.

## Results

### TDP-43 loss triggers cryptic exon inclusion and decreased synaptic and membrane excitability gene expression in human neurons and FTD-MND (motor neuron disease) patient brains

We previously re-analyzed an RNA-seq dataset in which neuronal nuclei with and without TDP-43 were sorted from FTD/ALS postmortem brain samples, and identified TDP-43-dependent cryptic splicing events in many synaptic genes (Liu et al., 2019; Ma et al., 2022). Because some of these cryptic splicing events could be caused either directly by TDP-43 reduction or by indirect effects of neurodegeneration, we performed experiments to individually validate these cryptic splicing events. We knocked down TDP-43 in human neurons derived from embryonic stem cells (iNeurons) using lentiviral shRNAs and examined individual cryptic splicing events using quantitative reverse transcription polymerase chain reaction (RT-qPCR). We tested two independent shRNAs targeting TDP-43, which produced similar results, and selected one for subsequent experiments. To minimize the possibility of secondary effects, such as stress responses, we focused on *bona fide* cryptic splicing events rather than those annotated as alternative splicing in the reference genome. We also excluded splicing changes that were difficult to detect by RT-qPCR (e.g., only one side of splicing junction was captured in these events). Following TDP-43 knockdown for seven days, we confirmed cryptic splicing changes in ten additional genes beyond *STMN2* and *UNC13A: AKT3*, *CACNA1E*, *CEP290*, *KALRN*, *KCNQ2*, *MNAT1*, *RAP1GAP*, *SETD5*, *STXBP5L*, and *SYT7* (Figure 1A). These splicing events exhibited effect sizes ranging from 3-to 50-fold with various types of splicing changes, including cryptic exon inclusion (*AKT3*, *CEP290*, *KALRN*, *MNAT1*, *RAP1GAP*, *SYT7*), 5’ splice site shift (*CACNA1E*, *SETD5*, *STXBP5L*), and exon skipping (*KCNQ2*).

**Figure 1.**
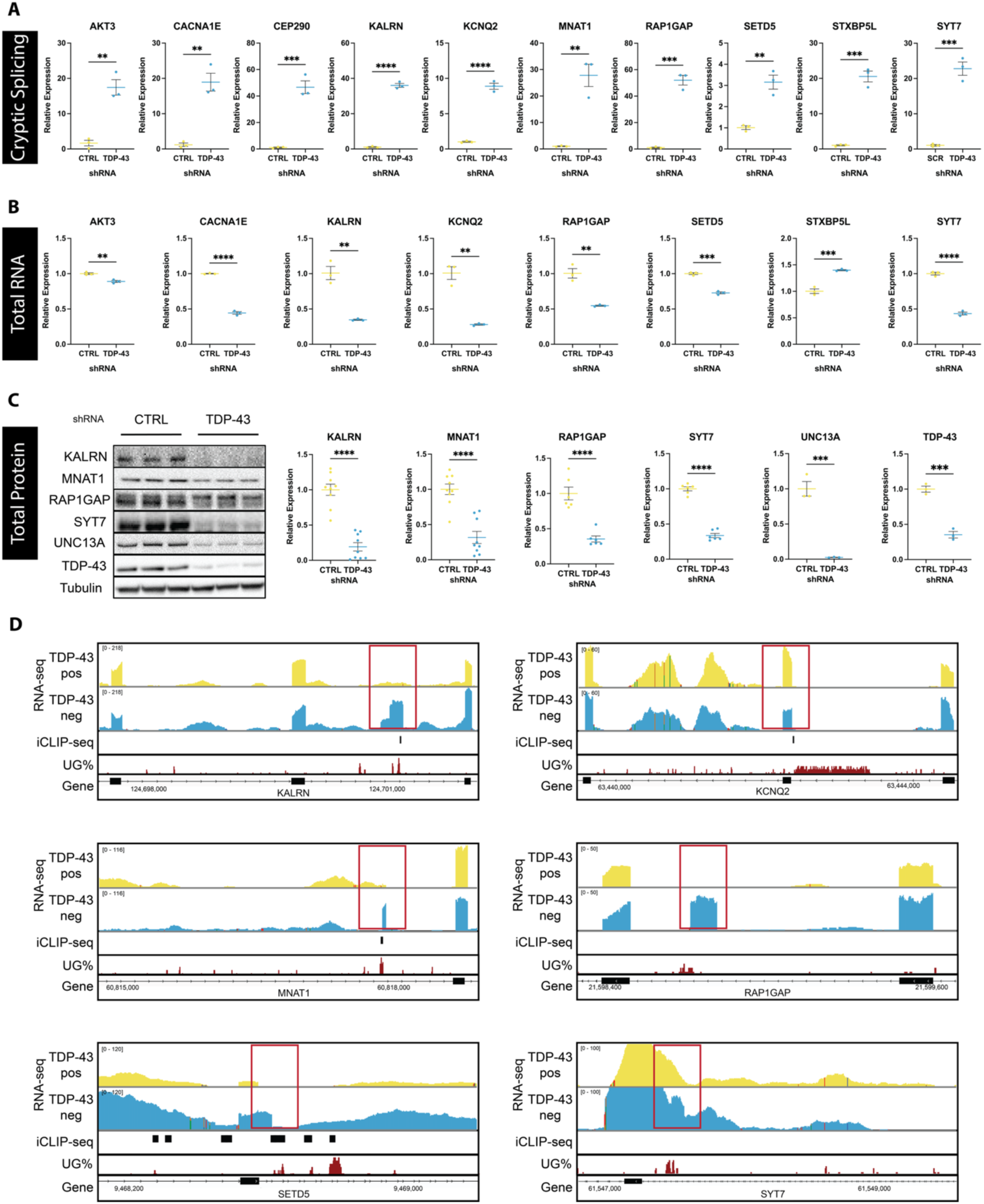
TDP-43 loss of function directly leads to cryptic splicing and reduced gene expression A. The levels of transcripts with cryptic exons in *AKT3*, *CACNA1E*, *CEP290*, *KALRN*, *KCNQ2*, *MNAT1*, *RAP1GAP*, *SETD5*, *STXBP5L*, and *SYT7* were increased upon TDP-43 knockdown. qPCR experiments were performed in iNeurons treated with TDP-43 shRNAs for seven days. The level of *GAPDH* was used for normalization. The level in control condition was set to 1. Mean ± s.e.m., n=3, unpaired t-test, ** p<0.01, *** p<0.001, **** p<0.0001. B. The total transcript levels of *AKT3*, *CACNA1E*, *KALRN*, *KCNQ2*, *RAP1GAP*, *SETD5*, and *SYT7* were decreased upon TDP-43 knockdown while the total transcript level of *STXBP5L* was increased. qPCR experiments were performed in iNeurons treated with TDP-43 shRNAs for seven days. The level of *GAPDH* was used for normalization. The level in control condition was set to 1. Mean ± s.e.m., n=3, unpaired t-test, ** p<0.01, *** p<0.001, **** p<0.0001. C. The total protein levels of *UNC13A*, *KALRN*, *MNAT1*, *RAP1GAP*, and *SYT7* were decreased upon TDP-43 knockdown. Western blotting experiments were performed in iNeurons treated with TDP-43 shRNAs for twelve days. The level of GAPDH or Tubulin was used for normalization. The level in control condition was set to 1. Mean ± s.e.m., n=3, unpaired t-test, *** p<0.001, **** p<0.0001. D. TDP-43 binding sites or UG-rich motifs were detected in the cryptic splicing region of *KALRN*, *KCNQ2*, *MNAT1*, *RAP1GAP*, *SETD5*, and *SYT7*. Lane 1, RNA-seq track of TDP-43 positive nuclei from FTD/ALS patient brain tissues; Lane 2, RNA-seq track of TDP-43 negative nuclei from FTD/ALS patient brain tissues; Lane 3, CLIP-seq track of TDP-43 in SH-SY5Y cells; Lane 4, percentage of UG (TG) or GU (GT) dinucleotides in 20bp bins, displayed as IGV tracks with a data range of 0.4–1; red rectangle, cryptic exon.

Each TDP-43-regulated cryptic splicing event showed different sensitivity to TDP-43 levels. Knocking down TDP-43 in iNeurons for 3, 5, or 7 days, lowered TDP-43 levels to 27%, 15%, and 11% of its levels under control conditions, respectively (Figure S1A). Interestingly, some cryptic splicing events (*KALRN*, *CACNA1E*, *MNAT1*) appeared early, on Day 3 following TDP-43 knockdown and their levels steadily increased over time, whereas some other ones (*UNC13A*, *SYT7*, *SETD5*, *STXBP5L*, *CEP290*) did not appear until Day 5 or 7 (Figure S1B, C). Some cryptic targets are more sensitive to TDP-43 loss than others and therefore the level of cryptic splicing did not always positively correlate with the reduced level of TDP-43, such as in the case of *STMN2*, *RAP1GAP*, *KCNQ2*, and *AKT3* (Figure S1D).

Most of these cryptic splicing events occur in the coding region of genes and result in frameshifts to open reading frames, introducing premature termination codons (PTCs). The only two exceptions are *STXBP5L*, in which the cryptic splicing is in the 5’ untranslated region (5’ UTR), and *KCNQ2*, in which exon skipping does not lead to an open reading frame shift. Given that transcripts with PTCs can be degraded via nonsense-mediated RNA decay (NMD) (Kurosaki et al., 2019; Lykke-Andersen and Jensen, 2015) and it has been shown that many cryptic splicing targets of TDP-43 are NMD substrates (Sinha et al., 2025; Zeng et al., 2025), does the accumulation of cryptic splicing transcripts with PTCs trigger NMD activation and subsequently decrease normal gene expression of these targets? Indeed, the level of total transcripts or normally spliced transcripts in multiple targets decreased significantly upon TDP-43 reduction (Figure 1B, Figure S1E). Consequently, following 12 days of TDP-43 knockdown, we found that the protein levels of multiple genes – *KALRN*, *MNAT1*, *RAP1GAP*, and *SYT7* – were reduced (Figure 1C). Therefore, in addition to *UNC13A* and *STMN2*, we have now identified additional TDP-43 targets that undergo cryptic splicing and show reduced gene expression in neurons upon TDP-43 reduction. To determine whether TDP-43 directly binds to these cryptic splicing targets, we analyzed publicly available crosslinking and immunoprecipitation sequencing (CLIP-seq) data (Zhao et al., 2022). This analysis revealed TDP-43 binding sites near cryptic splicing regions in *AKT3*, *CEP290*, *KALRN*, *KCNQ2*, *MNAT1*, *SETD5*, and *STXBP5L* (Figure 1D, Figure S1F). Because TDP-43 preferentially binds UG-rich motifs (Polymenidou et al., 2011; Tollervey et al., 2011), we quantified the local frequency of UG (TG) and GU (GT) dinucleotides and observed a higher percentage of UG-rich sequences within cryptic splicing regions of *AKT3*, *CEP290*, *KALRN*, *KCNQ2*, *MNAT1*, *RAP1GAP*, *SETD5*, *STXBP5L* and *SYT7* (Figure 1D, Figure S1F). These findings provide evidence that TDP-43 can directly bind and regulate these cryptic splicing events.

To confirm that these new cryptic splicing targets are associated with TDP-43 pathology in human brain, we performed RT-qPCR to detect individual cryptic splicing events in bulk postmortem brain tissues from FTD-MND patients. We observed increased cryptic splicing levels in the frontal cortex of patients, which is more vulnerable to TDP-43 pathology, whereas we did not in the cerebellum, which is less vulnerable (Figure S2A).

To directly visualize these splicing changes in patient tissues and relate cryptic splicing with TDP-43 pathology at single-cell resolution, we designed custom BaseScope *in situ* hybridization probes that specifically bind to the cryptic splicing junction of candidate TDP-43 targets, including *KALRN*, *KCNQ2*, *RAP1GAP*, *SETD5*, and *SYT7* (the sequences near the cryptic splicing junction of the other targets were not amenable to probe design). Combining *in situ* hybridization for individual cryptic splicing event with immunofluorescence for NeuN (to detect neuronal nuclei) and TDP-43 (to detect TDP-43 pathology), we stained frontal cortex sections from three FTD-MND patients with 3-6 sections per case per target. Across cortical Layer 2, we randomly sampled 1678, 1731, and 2165 NeuN+ neurons from each patient, of which 9.6% (±0.9%), 22.2% (±1.4%), and 33.9% (±1.5%) exhibited depletion of TDP-43 from the nucleus, respectively. In these neurons lacking nuclear TDP-43, we observed an accumulation of the cryptic splicing transcripts, with 58%, 81%, 83%, 63%, and 14% showing cryptic splicing of *KALRN*, *KCNQ2*, *RAP1GAP*, *SETD5*, and *SYT7*, respectively. By contrast, neighboring neurons without TDP-43 cytoplasmic inclusions rarely exhibited such signals (1.7%, 2.2%, 1.4%, 0.7%, and 0.8%, respectively) (Figure 2; Figure S2B, C). The extremely low number of puncta detected in TDP-43–positive neuronal nuclei may reflect background noise or transcripts originating from nearby TDP-43–negative cells in adjacent focal planes. We did not detect cryptic splicing transcripts in control brain tissue, further demonstrating that these are specific pathological events caused by loss of nuclear TDP-43 (Figure S3).

**Figure 2.**
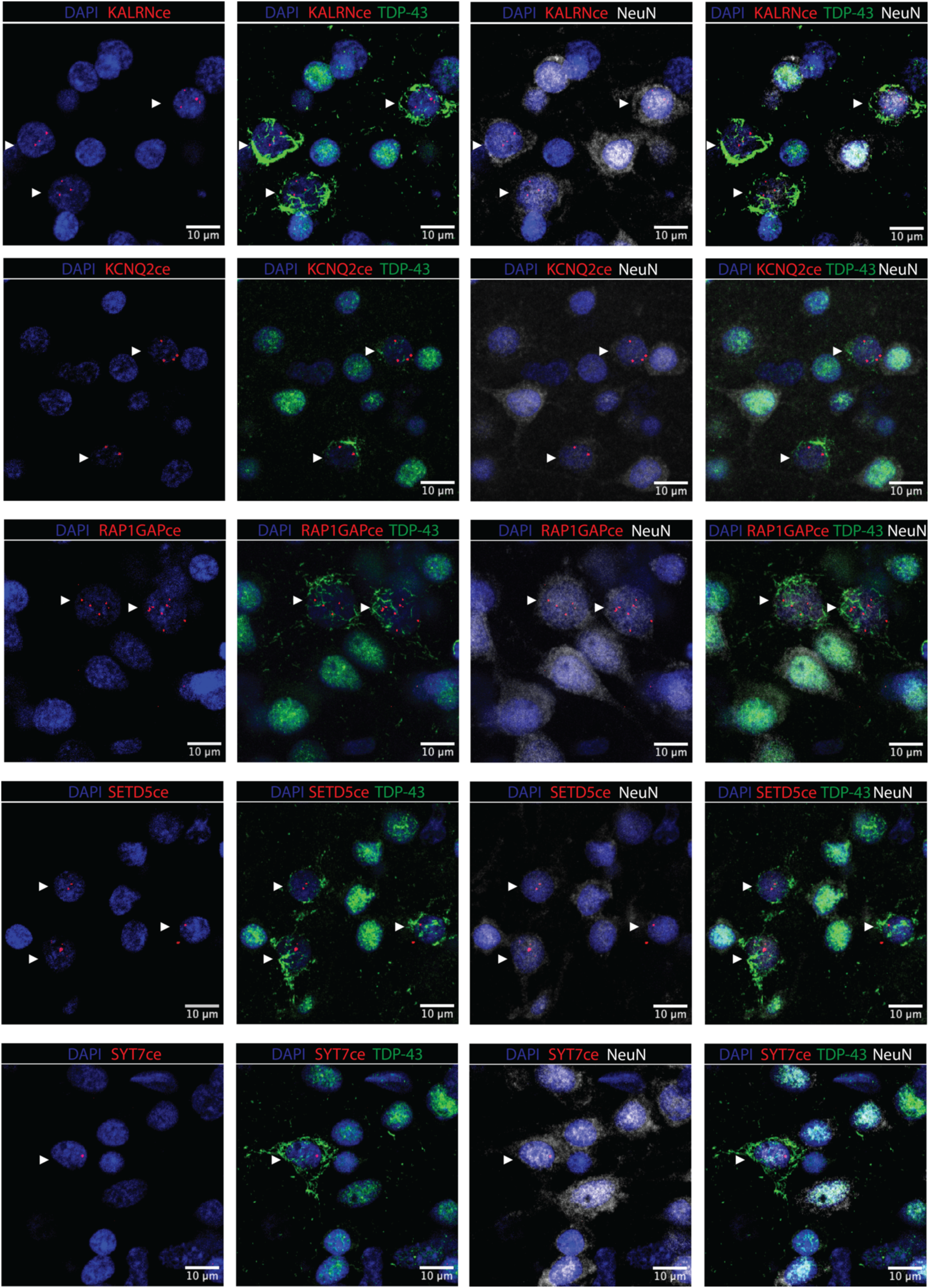
Cryptic splicing is present in neurons lacking nuclear TDP-43 in brains from FTD-MND patients BaseScope *in situ* hybridization shows puncta of transcripts with a cryptic exon in *KALRN*, *KCNQ2*, *RAP1GAP*, *SETD5*, and *SYT7* (red), in combination with immunofluorescent staining of the nuclear marker DAPI (blue), TDP-43 (green), and the neuronal marker NeuN (white) on sections from the medial frontal cortex of FTD-MND patients. Arrowheads indicate neurons lacking nuclear TDP-43 and with cytoplasmic TDP-43 aggregates.

Because we observed reduced expression of several cryptic splicing targets in our iNeuron analyses (Figure 1B), we designed BaseScope probes to detect the normally spliced transcripts to quantify gene expression levels. In control brain samples, these transcripts were readily detectable in neurons, whereas their levels were markedly decreased in patient brain samples harboring TDP-43 pathology (Figure S2D). Even within patient samples, those neurons with TDP-43 pathology showed a dramatic reduction in normally spliced transcripts compared to neighboring neurons with normal nuclear TDP-43 (Figure S4). Taken together, the histological observations confirms that cryptic splicing in *KALRN*, *KCNQ2*, *RAP1GAP*, *SETD5* and *SYT7* is strongly associated with nuclear TDP-43 depletion in the brains of patients with FTD-MND.

### Loss of TDP-43 or its cryptic splicing targets impairs synaptic function and alters membrane excitability

Several of these TDP-43 cryptic splicing targets encode proteins essential for neuronal function. Kalirin (KALRN) is a Rho-GTP exchange factor that regulates dendritic spine formation and synaptic plasticity (Parnell et al., 2021; Paskus et al., 2020). *Kalrn* knockout mice display an age-dependent reduction in spine density and glutamatergic transmission in the frontal cortex (Cahill et al., 2009). Mutations in *KALRN* have been linked to various neurological disorders such as autism spectrum disorder, schizophrenia, and intellectual disability (Parnell et al., 2021). RAP1 GTPase activating protein (RAP1GAP) is a negative regulator of Rap1 GTPase, which controls dendritic spine morphology and synaptic depression via the lysosome p38MAPK pathway (McAvoy et al., 2009; Zhang et al., 2018). Beyond postsynaptic signaling, TDP-43-dependent cryptic splicing also affects presynaptic genes like synaptotagmin 7 (SYT7), a member of the calcium sensor family which is important for synaptic vesicle release (Courtney et al., 2023; Luo and Südhof, 2017; Sugita et al., 2001; Vevea et al., 2021; Wu et al., 2024). *Syt7* knockout mice exhibit alterations in short-term synaptic plasticity (Jackman et al., 2016) and mutations in *SYT7* have been associated with bipolar disorder (Wang et al., 2020). We also identified a cryptic splicing event in the potassium voltage-gated channel subfamily Q member 2 (*KCNQ2*), which produces the Kv7.2 ion channel responsible for the neuronal M-current that regulates subthreshold membrane excitability (Wang et al., 1998). Pathogenic variations in *KCNQ2* are associated with benign familial neonatal seizures and developmental and epileptic encephalopathy (Biervert et al., 1998; Singh et al., 1998; Weckhuysen et al., 2012). Intrinsic membrane hyperexcitability has been observed in ALS patients and in patient iPSC-derived motor neurons carrying *superoxide dismutase* (*SOD1*), *C9orf72*, and *fused-in-sarcoma* (*FUS*) mutations. Pharmacological activation of Kv7 channels reduces this hyperexcitability and promotes motor neuron survival *in vitro*, and has recently been tested as a strategy to decrease cortical and spinal motor neuron excitability in ALS patients (Wainger et al., 2021, 2014). Finally, *MNAT1*, a component of the CDK-activating kinase (CAK) complex with cyclin H and CDK7, is required for transcription regulation (Adamczewski et al., 1996; Yee et al., 1995). Although the function of MNAT1 in neurons remains poorly explored, its partner CDK7 has been implicated in activity-dependent regulation of neuronal gene expression (He et al., 2017). Notably, a recent study identified the same *MNAT1* cryptic exon in human iPSC-derived motor neurons and ALS/FTD patient brain tissue as a result of TDP-43 loss (Tanaka et al., 2025). Taken together, these TDP-43 splicing targets are crucial for neuronal function and therefore their cryptic splicing and reduced expression could result in significant functional impairments.

Next, to define how TDP-43 and its splicing targets contribute to neuronal function, we individually knocked down each gene and monitored neuronal activity using MEA, which enables simultaneous measurement of neuronal spontaneous firing (indicated by weighted mean firing rate), excitability (indicated by number of bursts), and network connectivity (indicated by synchrony index). We co-cultured human iNeurons with mouse astrocytes on MEA plates with 16 electrodes in each well. By Day 21, neurons exhibited robust activity indicative of synaptic network formation, at which point we introduced lentiviral shRNAs targeting TDP-43 or its splicing targets – *KALRN, RAP1GAP, SYT7, UNC13A, MNAT1, KCNQ2*, and *STMN2* (Figure 3A). We performed the experiments in two separate batches, because we included a large number of replicates and only a limited number of conditions can be tested in each run. We observed batch-to-batch variability—likely due to factors such as electrode performance, infection efficiency, and cell density—but we analyzed and present the data for each batch separately and restrict comparisons within each batch to ensure robust conclusions.

**Figure 3.**
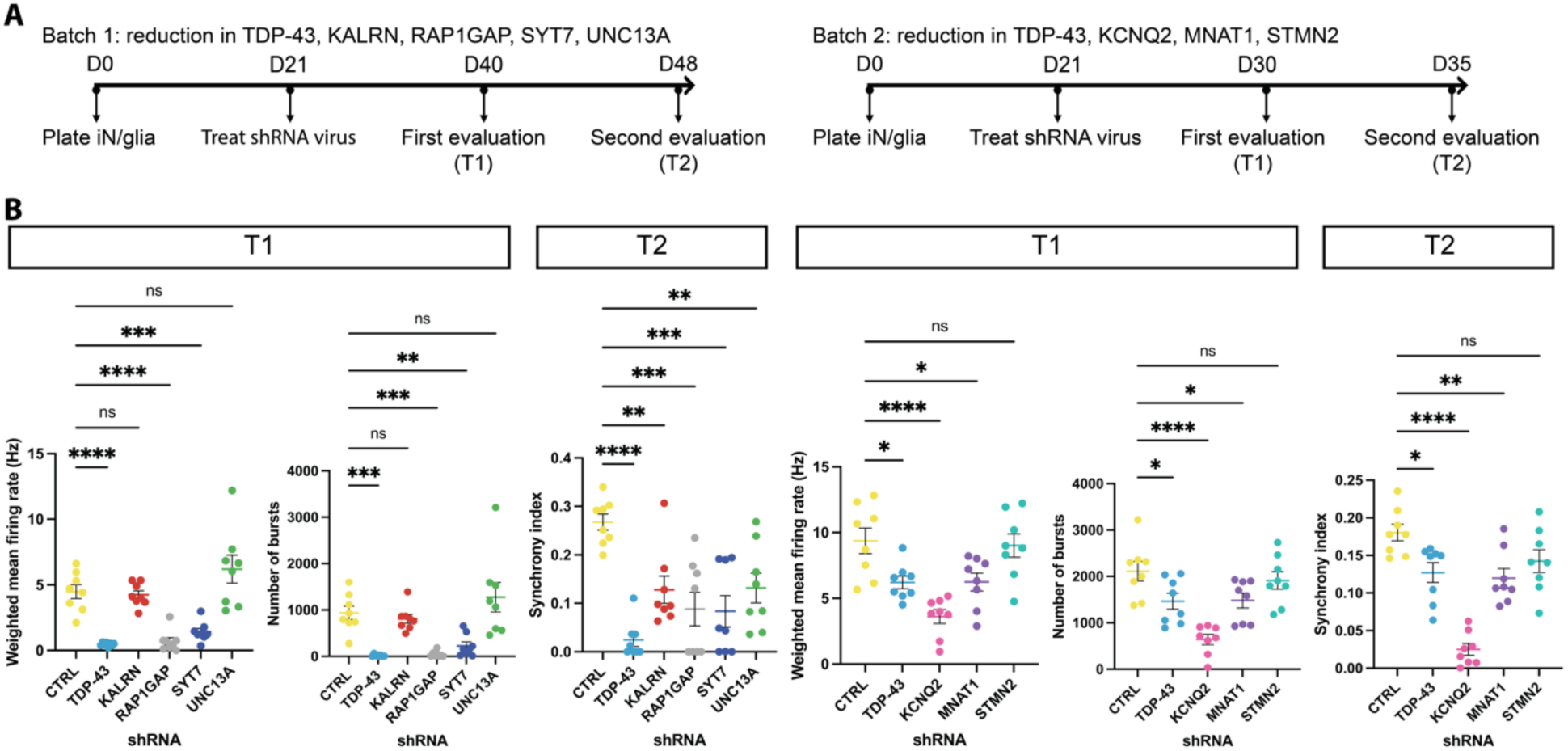
Cryptic splicing targets of TDP-43 are critical for maintaining neuronal activity A. Schematic diagrams show the experimental design of multielectrode array (MEA) recordings upon knockdown of TDP-43 or its cryptic splicing targets in iNeurons in two batches. Batch 1 examined the effects of a reduction in TDP-43, KALRN, RAP1GAP, SYT7, and UNC13A, and Batch 2 examined the effects of a reduction in TDP-43, KCNQ2, MNAT1, and STMN2. Knockdown or scramble shRNAs were administered on Day 21 in both batches. A first evaluation was conducted on Day 40 in Batch 1 and on Day 30 in Batch 2 (T1). A second evaluation was conducted on Day 48 in Batch 1 and on Day 35 in Batch 2 (T2). B. MEA analysis shows decreases in the weighted mean firing rate (spontaneous firing) and the number of bursts (excitability) upon reductions in TDP-43, RAP1GAP, SYT7, KCNQ2, and MNAT1 at T1, as well as decreases in the synchrony index (connectivity) upon reductions in TDP-43, KALRN, RAP1GAP, SYT7, UNC13A, KCNQ2, and MNAT1 at T2. Mean ± s.e.m., n=8, one-way ANOVA, * p<0.05, ** p<0.01, *** p<0.001, **** p<0.0001.

Because mouse astrocytes were present, we designed shRNAs to specifically target human transcripts, ensuring that the observed phenotypes reflected neuronal knockdown. Knockdown efficiency ranged from 50% to 98% in iNeurons after a 7-day treatment, but showed no effect in mouse primary cultures, except for *KCNQ2*, which shares a high sequence conservation between mouse and human (Figure S5A, B).

Measurements from the MEA assay are based on field potential changes of the collective population activity; therefore, to exclude potential effects of cell loss on neuronal activity, we quantified the number of covered electrodes, an MEA readout indicating culture integrity by measuring cell impedance. Most knockdowns – *KALRN*, *RAP1GAP*, *SYT7*, *UNC13A*, *MNAT1*, and *STMN2* – did not significantly reduce electrode coverage throughout the recording period (Day 48/T2 in batch 1, Day 35/T2 in batch 2). In contrast, TDP-43 knockdown and *KCNQ2* knockdown maintained the number of covered electrodes up until Day 40 (T1 in batch 1) and Day 30 (T1 in batch 2), respectively (Figure S5C). Based on these data, we identified a time window where the number of covered electrodes was comparable across conditions – so we could examine neuronal activity independent of cell loss.

TDP-43 knockdown significantly impaired spontaneous neuronal firing, excitability, and connectivity. The weighted mean firing rate and the number of bursts were markedly reduced within 7 days of knockdown, indicating a rapid functional decline (Figure 3B, Figure S5D-F). Similarly, knockdown of multiple TDP-43 splicing targets, including *RAP1GAP*, *SYT7*, *MNAT1*, and *KCNQ2*, robustly decreased neuronal activity across multiple MEA metrics reflecting altered excitability and connectivity (Figure 3B, Figure S5D-F). On the contrary, knockdown of *UNC13A* and *KALRN* did not significantly change spontaneous firing or excitability, but reduced network connectivity at a later time point (T2, Figure 3B, Figure S5D, E), consistent with their known functions in synaptic transmission. In addition to the synaptic genes, we also examined the TDP-43 cryptic splicing target *STMN2*, which is involved in axon growth and regeneration. As expected, *STMN2* knockdown showed minimal effect on synaptic activity or membrane excitability (Figure 3B, Figure S5D, F). Collectively, these data provide evidence that TDP-43 and its splicing targets are essential for maintaining membrane excitability and synaptic activity, and their loss of function profoundly impairs neuronal function.

### Blocking cryptic splicing rescues impaired neuronal activity caused by TDP-43 loss in human neurons

To test if cryptic splicing of these synaptic targets directly contribute to the neuronal dysfunction caused by TDP-43 loss, we developed antisense oligonucleotides (ASOs) – short synthetic single-stranded DNAs – that specifically bind and inhibit individual cryptic splicing events in *KALRN*, *RAP1GAP*, *SYT7*, and *UNC13A* (Figure 4A). For each gene target, we screened multiple ASO candidates targeting sequences at the cryptic splicing junctions or within the cryptic splicing exon in iNeurons with TDP-43 knockdown and selected the ones that most effectively suppressed the cryptic splicing event. We treated iNeurons with each ASO on Days 3 and 10 after plating with TDP-43 shRNAs administered on Day 5, and collected RNAs and proteins on Day 12 and Day 15, respectively. The ASOs effectively reduced cryptic splicing (Figure 4B) and, importantly, partially restored total RNA (Figure 4C) and protein expression (Figure 4D). The level of TDP-43 did not change upon ASO treatment, confirming that the inhibition of cryptic splicing is a direct result of ASO blockage (Figure S6A).

**Figure 4.**
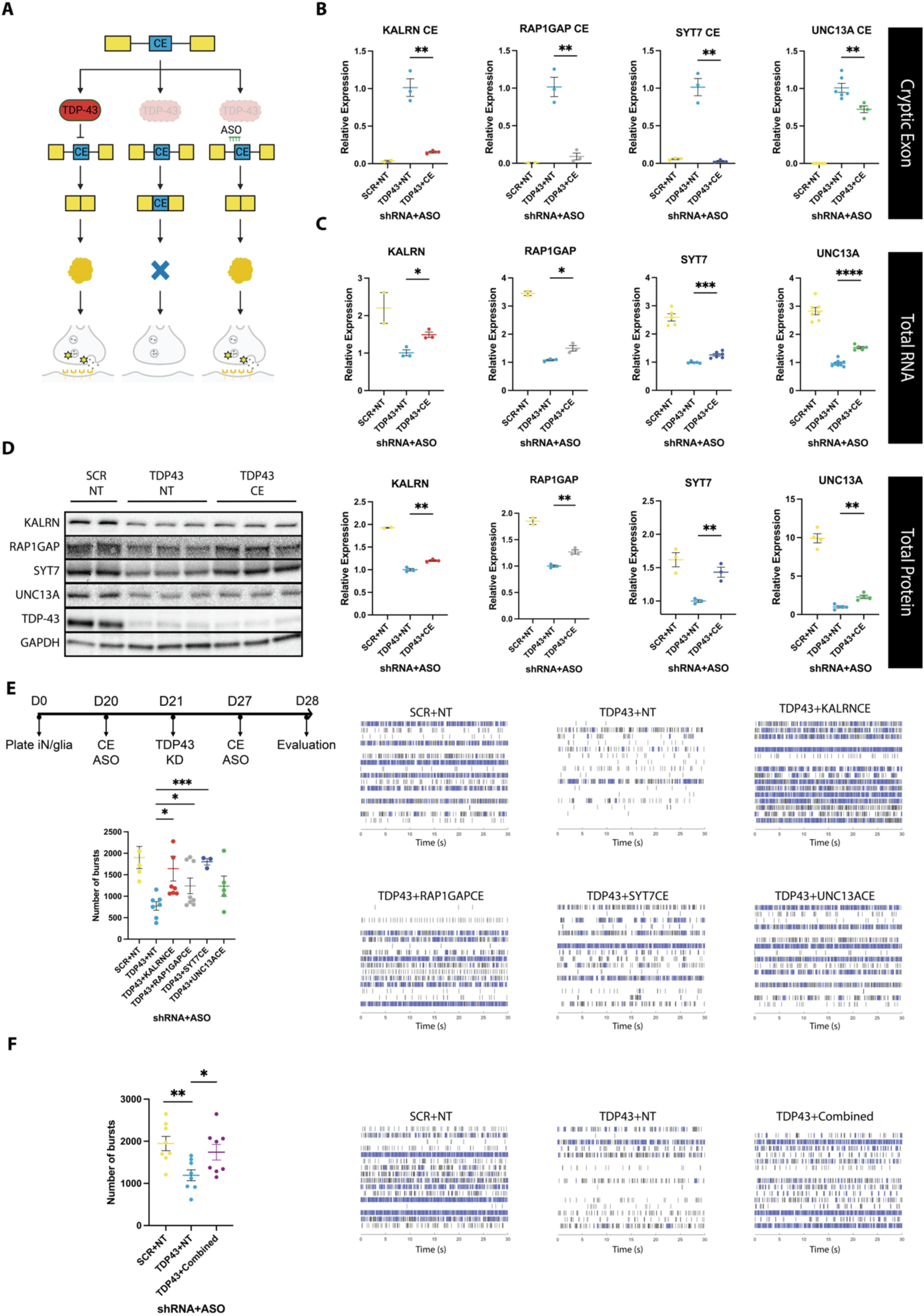
Inhibiting cryptic splicing of synaptic genes restores the neuronal dysfunction caused by TDP-43 loss A. Schematic diagram shows the model in which individual splicing-inhibiting ASOs can reduce the cryptic splicing level in an individual target, restore its normal gene expression, and eventually ameliorate neuronal dysfunction caused by TDP-43 loss. B. TDP-43 reduction increases the level of transcripts with cryptic exons in *KALRN*, *RAP1GAP*, *SYT7*, and *UNC13A*, which is inhibited by adding ASOs targeting each cryptic splicing event individually. qPCR experiments were performed in iNeurons treated with TDP-43 or scramble (SCR) shRNAs for 7 days and cryptic exon-blocking (CE) or non-targeting (NT) ASOs for 9 days. The level of *GAPDH* was used for normalization. The level in TDP-43 knockdown condition was set to 1. Mean ± s.e.m., n=2-6, unpaired t-test, ** p<0.01. C. TDP-43 reduction decreases the level of total transcripts in *KALRN*, *RAP1GAP*, *SYT7*, and *UNC13A*, which is partially reversed by adding ASOs targeting each cryptic splicing event individually. qPCR experiments were performed in iNeurons treated with TDP-43 or scramble (SCR) shRNAs for 7 days and cryptic exon-blocking (CE) or non-targeting (NT) ASOs for 9 days. The level of *GAPDH* was used for normalization. The level in TDP-43 knockdown condition was set to 1. Mean ± s.e.m., n=2-7, unpaired t-test, * p<0.05, *** p<0.001, **** p<0.0001. D. TDP-43 reduction decreases the level of total proteins in KALRN, RAP1GAP, SYT7, and UNC13A which is ameliorated by adding ASOs targeting each cryptic splicing event individually. Immunoblotting experiments were performed in iNeurons treated with TDP-43 or scramble (SCR) shRNAs for 10 days and cryptic exon-blocking (CE) or non-targeting (NT) ASOs for 12 days. Representative images are shown on the left and quantifications on the right. The level of GAPDH was used for normalization. The level in TDP-43 knockdown condition was set to 1. Mean ± s.e.m., n=2-4, unpaired t-test, ** p<0.01. E. TDP-43 reduction decreases the number of bursts, which is partially reversed by adding ASOs targeting cryptic splicing in *KALRN*, *RAP1GAP*, *SYT7*, and *UNC13A* individually. MEA analysis was conducted as shown by the schematic diagram on the top left: co-culture was treated with TDP-43 or scramble (SCR) shRNAs on Day 21 and cryptic exon-blocking (CE) or non-targeting (NT) ASOs on Days 20 and 27; evaluation was conducted on Day 28. Quantifications are shown on the bottom left, mean ± s.e.m., n=3-8, unpaired t-test, * p<0.05, *** p<0.001. Representative raster plots on the right show neuronal activity in a 30-second time frame for each condition. F. TDP-43 reduction decreases the number of bursts, which is reversed by adding a combinations of ASOs targeting cryptic splicing in *KALRN*, *RAP1GAP*, *SYT7*, and *UNC13A*. MEA analysis was conducted as shown by the schematic diagram in E. Quantifications are shown on the left, mean ± s.e.m., n=8, unpaired t-test, * p<0.05, ** p<0.01. Representative raster plots on the right show neuronal activity in a 30-second time frame for each condition.

We next investigated if inhibiting cryptic splicing is sufficient to rescue the neuronal activity impairments caused by loss of TDP-43. We treated iNeurons with each ASO twice on Days 20 and 27 after cell plating and TDP-43 shRNAs on Day 21 (Figure 4E). In neurons treated with non-targeting scramble shRNAs, ASO treatment did not significantly change neuronal activity (Figure S6B). However, upon TDP-43 reduction, each ASO partially mitigated the neuronal activity impairment prior to any cell loss (Figure 4E, Figure S6C). Intriguingly, administration of a combination of all four ASOs was sufficient to almost fully alleviate TDP-43-loss induced neuronal dysfunction (Figure 4F, Figure S6C). Together, these results provide evidence that each of these cryptic splicing events contributes, at least partially, to the neuronal dysfunction mediated by TDP-43 loss and provide a set of ASOs that effectively suppress the cryptic splicing of these neuronal TDP-43 targets.

## Discussion

In this study, we report additional cryptic splicing targets of TDP-43, beyond *UNC13A* and *STMN2*. Using human stem cell-derived neurons, we demonstrate that TDP-43 reduction directly causes abnormal splicing of these genes, several of which were also detected in iNeurons from ALS/FTD patients (Rothstein et al., 2025). Most of these new cryptic splicing events introduce premature termination codons, resulting in decreases in the level of their normal RNA and protein. Using CLIP-seq and UG-motif analysis, we found significant enrichment of TDP-43 binding sites in proximity to the cryptic splicing regions, further confirming the direct role of TDP-43 in these cryptic splicing events. Applying *in situ* hybridization to human postmortem brain tissues, we demonstrated that several cryptic splicing events can be detected selectively in neurons lacking nuclear TDP-43 in patients with FTD-MND, providing further evidence that these targets could contribute to disease pathogenesis. Because of methodological limitations in the number of cryptic splicing events that can be evaluated simultaneously, we were unable to assess whether these cryptic splicing events appear in the same neurons lacking nuclear TDP-43, or what subtypes of neurons they are present in. Interestingly, a recent single nucleus sequencing study revealed that the cryptic splicing events in *STMN2* and *KALRN* can be detected in frontal cortex excitatory neurons, especially the Layer 5 extratelencephalic projection neurons known to be selectively vulnerable in some forms of FTD (Hodge et al., 2020), in patients with FTD due to a *C9orf72* expansion (Gittings et al., 2023). In the future, multiplexed *in situ* hybridization or spatial transcriptomics should be utilized to further dissect the vulnerable neuronal subtypes affected by cryptic splicing.

We provide evidence that multiple cryptic splicing targets play critical roles in membrane excitability and synaptic function, including *UNC13A*, *SYT7*, *KALRN*, *RAP1GAP*, *MNAT1*, and *KCNQ2* – knocking down these genes or TDP-43 itself impaired neuronal activity. Synaptic dysfunction has been reported as an early symptom before neurodegeneration in ALS and FTD (Wilson et al., 2023). Several imaging and cerebrospinal fluid biomarkers that evaluate synaptic health have been applied to diagnosis and prognosis of ALS and FTD (Dejanovic et al., 2024). In our *in vitro* study of iNeurons, we found altered excitability and synaptic dysfunction precedes neuronal loss when knocking down TDP-43 or its cryptic splicing targets, further demonstrating that neuronal dysfunction caused by loss of function in membrane excitability and synaptic genes could serve as a critical factor during neurodegeneration.

An intriguing finding is that *KCNQ2* knockdown in iNeurons markedly decreases neuronal activity, which contrasts with previous reports that KCNQ2 inhibition leads to hyperexcitability (Wainger et al., 2021, 2014). One potential explanation is that, unlike acute M-current inhibition, chronic M-current depression may drive a distal shift of the axon initial segment and redistribution of Na^+^ and Kv7 channels, thereby raising the firing threshold and reducing intrinsic excitability (Lezmy et al., 2017). Alternatively, loss of KCNQ2 may induce compensatory upregulation of conductance via other K⁺ channels, paradoxically dampening excitability; indeed, faster action potential repolarization, larger post-burst afterhyperpolarization, and enhanced Ca²⁺-activated K⁺ currents have been reported in patient-derived iNeurons with *KCNQ2* mutations (Simkin et al., 2021). Finally, cell-type specific differences between NGN2-induced cortical neurons and motor neurons, or potential off-target effects of *KCNQ2* shRNA, may also underlie the reduced firing phenotype.

To determine whether cryptic splicing in these synaptic genes directly contributes to the neuronal dysfunction caused by loss of TDP-43, we developed splice-modulating ASOs that specifically inhibit the cryptic splicing events in *UNC13A, SYT7, KALRN*, and *RAP1GAP*. Treating neurons with these ASOs effectively decreased cryptic splicing transcripts caused by TDP-43 loss.

The effect on restoring total RNA and protein levels was not complete, likely owing to the short time period of these *in vitro* experiments. Nevertheless, treating neurons with these ASOs was sufficient to mitigate the impairment in neuronal activity caused by TDP-43 loss. ASOs targeting single cryptic splicing events produce partial rescue, with the ASO blocking the *KALRN* cryptic exon showing the strongest effect. A combination treatment of ASOs targeting four TDP-43 targets at once resulted in near-complete restoration of activity. These results underscore the broad impact of TDP-43 loss on various synaptic and membrane excitability genes and suggest that therapies aimed at a single cryptic splicing event may not be as efficacious as targeting multiple ones simultaneously (Wilkins et al., 2024). In summary, we present multiple membrane excitability and synaptic genes as cryptic splicing targets of TDP-43, whose misregulation drives neuronal dysfunction. Suppressing cryptic splicing of these genes, individually or in combination, is sufficient to mitigate the neuronal activity deficits caused by loss of TDP-43. These findings provide insights and new targets for developing therapeutic strategies to combat TDP-43 pathology in ALS/FTD.

## Acknowledgments

This work was supported by NIH grants R35NS137159, U54NS123743, R01AG064690 (A.D.G.), Target ALS (A.D.G.), the Robert Packard Center for ALS Research at Johns Hopkins (A.D.G.), Milton Safenowitz Postdoctoral Fellowship from ALS Association (C.G.), and Dr. Miriam and Sheldon G Adelson Medical Research Foundation (K.C.). The UCSF Neurodegenerative Disease Brain Bank is supported by NIH grants AG019724, AG062422, AG063911, and AG057195; the Rainwater Charitable Foundation, and the Bluefield Project to Cure FTD. A.D.G. is a Chan Zuckerberg Biohub – San Francisco Investigator. We thank Stanford Gene Vector and Virus Core (GVVC) for lentiviral production.

## Methods

### Stem cell culture

Human H1 embryonic stem cells (hESCs) were maintained in mTeSR plus media (StemCell Technologies, 100-0276) on plates coated with Matrigel (Corning, 354230), and passaged every 3-4 days using ReLeSR (StemCell Technologies, 100-0483). hESCs were differentiated into cortical-like neurons (iNeurons) by overexpressing transcription factor neurogenin 2 (NGN2) driven by a Tet-On induction system in DMEM/F12 media supplemented with 1 ug/mL doxycycline, as previously described (Bieri et al., 2019). Cells were dissociated using Accutase (StemCell Technologies, 07920) on day 3 of differentiation and replated on plates coated with poly-D-lysine (Thermo Scientific, A3890401) and mouse laminin (Sigma-Aldrich, L2020-1MG) in Neurobasal Medium (Thermo Fisher, 21103049) containing neurotrophic factors, BDNF and GDNF (R&D Systems). After four days of seeding, iNeurons were infected with shRNA lentivirus.

### shRNA and lentiviral packaging

shRNA sequences were listed in Supplementary Table 1. Lentiviral packaging was achieved by transfecting HEK293T cells with each shRNA expression vector (Sigma-Aldrich, MISSION shRNA) and third generation packaging vectors pRSV-REV, pMDLg/pRRE, pMD2.G(VSV-G) (Addgene), followed by supernatant collection and concentration using Lenti-X Concentrator (Takara, 631232). Viral titers were determined by serial dilution followed by puromycin viability test.

### RNA extraction and RT-qPCR

Total RNA from iNeurons was extracted using RNeasy Plus Micro Kit (Qiagen, 74034) per manufacturer’s instructions. Total RNA from human brain tissues was extracted using Trizol (Invitrogen, 15596018) followed by TURBO DNase treatment (Thermo Fisher Scientific, AM2238) and cleanup using RNeasy Mini Kit (Qiagen, 74106). Reverse transcription into cDNA was conducted by High Capacity cDNA Reverse Transcription Kit (Thermo Fisher Scientific, 43-688-13). RT-qPCR was performed using Applied Biosystems PowerUp SYBR Green Master Mix (Thermo Fisher Scientific, A25742) on a QuantStudio 3 Real-Time PCR System (Thermo Fisher Scientific). The primers used are listed in Supplementary Table 2.

### Immunoblotting

Cells were lysed in ice-cold RIPA buffer (Sigma-Aldrich, R0278) supplemented with Halt Protease Inhibitor Cocktail (Thermo Fisher, 78429) on ice for 30 minutes with vortexing every 10 minutes, followed by centrifuging at 15,000 rpm for 15 minutes at 4 °C. The supernatant was collected and BCA protein assay (Invitrogen, 23225) was used to quantify protein concentrations. 10-20ug of protein of each sample was denatured in LDS sample buffer (Thermo Scientific, B0007) containing 5% 2-mercaptoethanol (Sigma-Aldrich) at 70 °C for 10 minutes. Samples were loaded to NuPAGE 4–12% Bis-Tris Protein Gels (Thermo Fisher, NP0335BOX) for gel electrophoresis, transferred onto 0.2 µm PVDF membranes (Bio-Rad, 162-0174) in NuPAGE Transfer Buffer (Life Technologies, NP0006-1) at 100 V for 2 h at 4 °C. Membranes were blocked in EveryBlot Blocking Buffer (Bio-Rad, 12010020), incubated using primary antibodies at 4 °C overnight, washed using Tris-buffered saline with 0.1% Tween-20 (TBST), and incubated with HRP-conjugated secondary antibodies (anti-mouse or anti-rabbit 1:10000, Thermo Scientific) at room temperature for 2 hours, and washed in TBST. Blots were detected by ECL Western Blotting Detection Reagent (Cytiva, RPN2106) and imaged using the ChemiDoc XRS+ System (Bio-Rad). The intensity of bands was quantified using Fiji. Primary antibodies used in this study were: TDP-43 (Proteintech, 10782-2-AP), Tubulin (Cell Signaling Technology, 2144S), GAPDH (Sigma-Aldrich, G8795), UNC13A (Proteintech, 68483-1-Ig), KALRN (Proteintech, 19740-1-AP), RAP1GAP (Abcam, ab32373), SYT7 (Thermo Scientific, PA5-52998), MNAT1 (Proteintech, 11719-1-AP).

### Human subjects and diagnostic neuropathological assessment

Postmortem brain tissue samples used for this study were obtained from the University of California San Francisco (UCSF) Neurodegenerative Disease Brain Bank. Supplementary Table 3 provides demographic, clinical, and neuropathological information. Consent for brain donation was obtained from subjects or their surrogate decision makers in accordance to the Declaration of Helsinki, and following a procedure approved by the UCSF Committee on Human Research. Clinical and neuropathological diagnosis were rendered as described previously (Nana et al., 2019). Subjects were selected based on clinical and neuropathological assessment. Patients selected had a primary clinical diagnosis of behavioral variant frontotemporal dementia (bvFTD) with or without amyotrophic lateral sclerosis (ALS)/motor neuron disease (MND) and a neuropathological diagnosis of frontotemporal lobar degeneration (FTLD)-TDP, Type B with or without ALS or MND. We excluded subjects if they had a known disease-causing mutation, post-mortem interval ≥ 24 h, Alzheimer’s disease neuropathologic change (ADNC) > low, Thal amyloid phase > 2, Braak neurofibrillary tangle stage > 4, CERAD neuritic plaque density > sparse, and Lewy body disease > brainstem predominant (Nana et al., 2019).

### *In situ* hybridization and immunofluorescence

Brains were cut fresh into ∼1 cm thick coronal slabs and alternating slabs were fixed in 10% neutral buffered formalin for 72 h or rapidly frozen. Blocks from medial frontal pole were dissected from the fixed coronal slabs, cryoprotected in graded sucrose solutions, frozen, and cut into 50 µm thick sections as described previously (Nana et al., 2019). To detect single RNA molecules, a BaseScope Red Assay kit (ACDBIO, USA) was used. For each cryptic splicing target, probes targeting the cryptic splicing junction or the normal splicing junction were used (sequence details in Supplementary Table 4). Positive (Homo sapiens PPIB) and negative (Escherichia coli DapB) control probes were also included. *In situ* hybridization was performed based on vendor specifications for the BaseScope Red Assay kit. Briefly, frozen tissue sections were washed in PBS and placed under an LED grow light (HTG Supply, LED-6B240) chamber for 48 h at 4 °C to quench tissue autofluorescence. Sections were quickly rinsed in PBS and blocked for endogenous peroxidase activity. Sections were transferred on to slides and dried overnight. Slides were subjected to target retrieval and protease treatment and advanced to *in situ* hybridization. Probes were detected with TSA Plus-Cy3 (Akoya Biosciences) and subjected to immunofluorescence staining with antibodies to TDP-43 (Proteintech, 10782-2-AP) and NeuN (Synaptic systems, 266004) and counterstained with DAPI (Life Technologies) for nuclei.

### Image acquisition and analysis

Z-stack images were captured using a Leica SP8 confocal microscope with an 63x oil immersion objective (1.4 NA). For RNA probes, image capture settings were established during initial acquisition based on PPIB and DAPB signal and remained constant across probes and subjects. TDP-43 and NeuN image capture settings were modified based on staining intensity differences between cases. For each case, 3 to 6 non-overlapping Z-stack images were captured across cortical layers 2-3. Fields of view (FOVs) were evenly distributed and selected solely based on DAPI staining. Each FOV contained an average of 76 NeuN+ neurons. For Figure S3B, we manually quantified the percentage of CE-positive neuronal nuclei with or without nuclear TDP-43, with each data point representing one FOV. For Figure S3C, we manually quantified the number of puncta per neuronal nucleus lacking nuclear TDP-43. For Figure S3D, we quantified the number of puncta per FOV in patient and control samples using FIJI.

### Multielectrode array

Maestro Pro multi-well MEA system (Axion Biosystems Inc., Atlanta, GA, USA) was used to perform MEA recording, with each well of 48-well MEA plates embedded with an array of 16 gold electrodes. Human iPSCs, LiPSC-GR1.1 (Lonza), were differentiated into iNeurons by overexpressing transcription factor neurogenin 2 (NGN2) driven by a Tet-On induction system. 11,250 mouse astrocytes and 75,000 iNeurons were spot seeded in the recording electrode area of each well in the Cytoview 48-well MEA plates. Half of the medium was changed every 3 to 4 days. Neuronal activity was recorded for 5 minutes using Axion BioSystems’ Integrated Studio software (AxIS, version 3.7.2) (Axion BioSystems) with a Butterworth band-pass filter of 10Hz and 2.5 kHz cutoff frequencies. Data analysis was performed by Neural Metric Tool (Axion Biosystems). A spike detection was computed with an adaptive threshold of six standard deviations of the estimated noise for each electrode. The electrodes that detected at least 5 spikes per minute were classified as active electrodes. Bursts were identified using an inter-spike interval (ISI) threshold requiring a minimum number of five spikes with a maximum ISI of 100ms. Network bursts were identified as bursts of >50 spikes that occurred in >35% of the active electrodes in the well, with a maximum ISI of 100ms. Synchrony represented a measure of similarity between two spike trains and the synchrony index was estimated by the area under the normalized cross-correlogram for a time window of 20 ms.

### ASO treatment

ASOs used in this study were synthesized by IDT. The sequences are available upon request. Each base is 2’-O-methoxyethyl-modified and every internucleotide linkage is a phosphorothioate, all cytosine residues are 5-methylcytosines. For iNeuron treatment, ASOs were added to the culture media for free uptake at 1 uM.

### Statistical tests

Statistics were calculated using Prism software by GraphPad. Individual data points are independent biological replicates. All error bars are SEM. Statistical significance was determined using 2-tailed unpaired Student’s t-tests for 2 group comparisons or one-way ANOVA with multiple comparisons for 3 or more comparisons unless otherwise indicated. ****, p <0.0001; ***, p < 0.001; **, p < 0.01; *, p <0.05; ns, p > 0.05.

## Competing Interests

A.D.G. is a scientific founder of Maze Therapeutics, Trace Neuroscience, and Lyterian Therapeutics. S.M., R.D.M., G.M., and E.M.G. are employees of Trace Neuroscience. Stanford University has filed a provisional patent on ASOs targeting some of the cryptic splicing events described in this manuscript.

---

**Figure S1.**
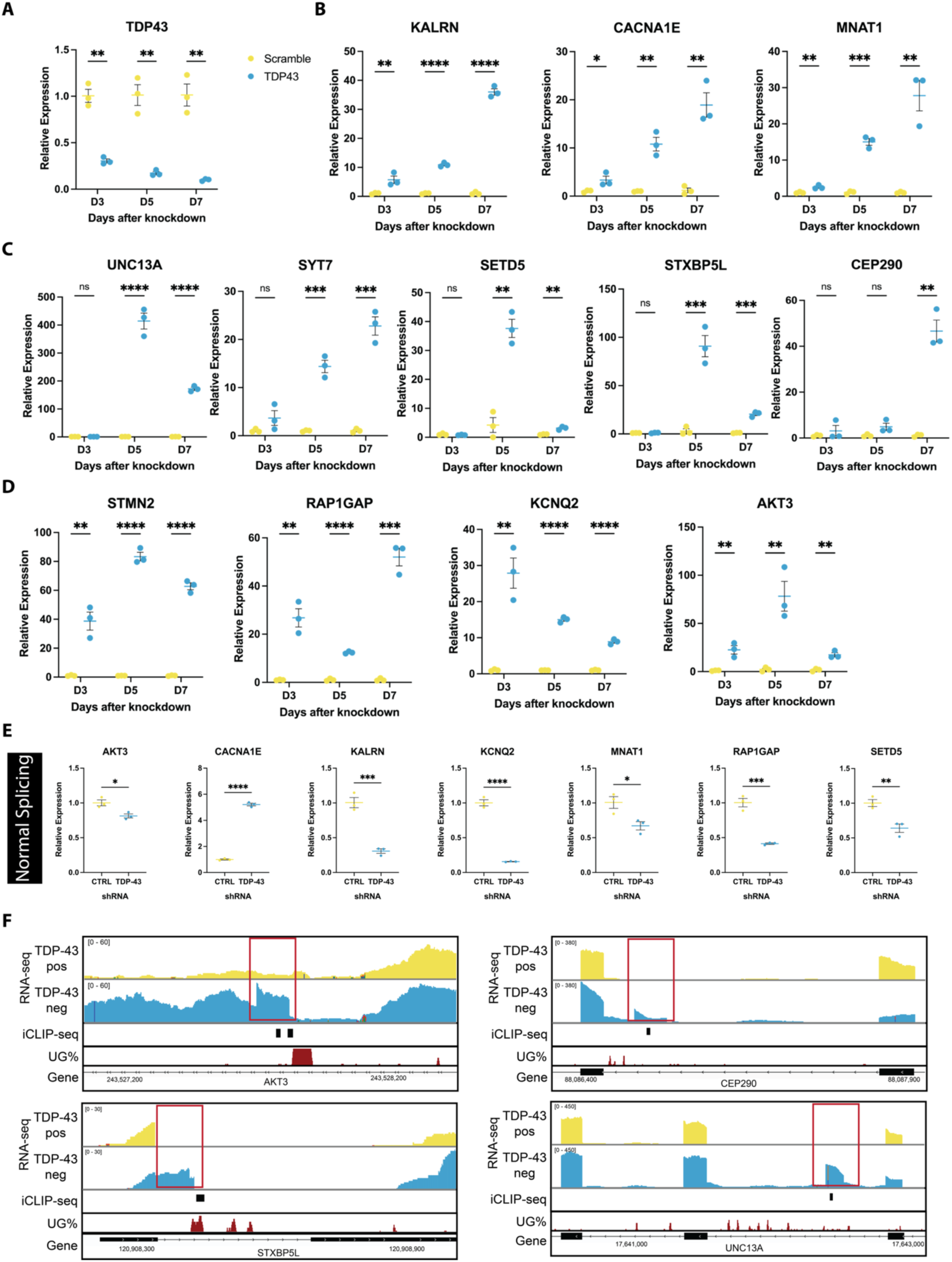
The level of cryptic splicing is sensitive to the reduced level of TDP-43 A. RT-qPCR shows decreased levels of TDP-43 transcripts when iNeurons treated with TDP-43 shRNAs for 3, 5, or 7 days. The level of *GAPDH* was used for normalization. The level in control condition at each time point was set to 1. Mean ± s.e.m., n=3, unpaired t-test, ** p<0.01. B. RT-qPCR shows the levels of cryptic splicing in *KALRN*, *CACNA1E*, and *MNAT1* can be detected 3 days after TDP-43 knockdown in iNeurons. The level of *GAPDH* was used for normalization. The level in control condition at each time point was set to 1. Mean ± s.e.m., n=3, unpaired t-test, * p<0.05, ** p<0.01, *** p<0.001, **** p<0.0001. C. RT-qPCR shows the levels of cryptic splicing in *UNC13A*, *SYT7*, *SETD5*, *STXBP5L*, and *CEP290* cannot be detected until 5 days after TDP-43 knockdown in iNeurons. The level of *GAPDH* was used for normalization. The level in control condition at each time point was set to 1. Mean ± s.e.m., n=3, unpaired t-test, ns not significant, ** p<0.01, *** p<0.001, **** p<0.0001. D. RT-qPCR shows the levels of cryptic splicing in *STMN2*, *RAP1GAP*, *KCNQ2*, and *AKT3* do not correlate with the level of TDP-43 in iNeurons. The level of *GAPDH* was used for normalization. The level in control condition at each time point was set to 1. Mean ± s.e.m., n=3, unpaired t-test, ** p<0.01, **** p<0.0001. E. RT-qPCR shows decreased levels of normal-spliced transcripts of *AKT3*, *CACNA1E*, *KALRN*, *KCNQ2*, *MNAT1*, *RAP1GAP*, and *SETD5* when iNeurons treated with TDP-43 shRNAs for seven days. The level of *GAPDH* was used for normalization. The level in control condition was set to 1. Mean ± s.e.m., n=3, unpaired t-test, * p<0.05, ** p<0.01, *** p<0.001, **** p<0.0001. F. TDP-43 binding sites or UG-rich motifs were detected in the cryptic splicing region of *AKT3*, *CEP290*, *STXBP5L*, and *UNC13A*. Lane 1, RNA-seq track of TDP-43 positive nuclei from FTD/ALS patient brain tissues; Lane 2, RNA-seq track of TDP-43 negative nuclei from FTD/ALS patient brain tissues; Lane 3, CLIP-seq track of TDP-43 in SH-SY5Y cells; Lane 4, percentage of UG (TG) or GU (GT) dinucleotides in 20bp bins, displayed as IGV tracks with a data range of 0.4–1; red rectangle, cryptic exon.

**Figure S2.**
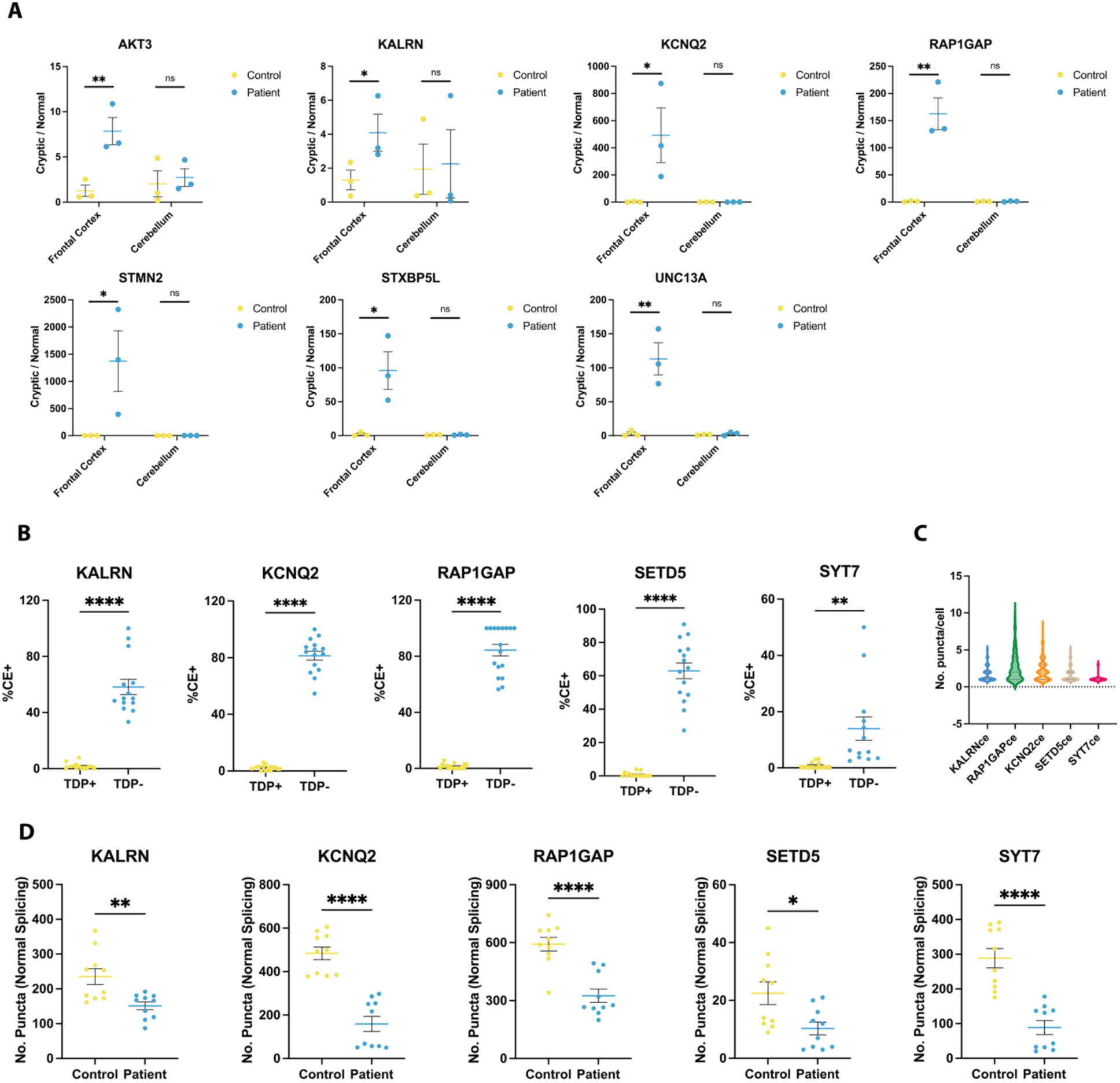
Cryptic splicing can be detected in postmortem brain tissues from FTD-MND patients A. RT-qPCR shows increased cryptic splicing in the frontal cortex but not in the cerebellum of patients. The level of normal-spliced transcripts was used for normalization. The level in control condition was set to 1. B. The percentage of neuronal nuclei that are positive for cryptic splicing puncta of each target in normal neurons (TDP+) or neurons with TDP-43 pathology (TDP-) according to BaseScope staining in patients with FTD-MND. Quantifications are based on 4 to 6 z-stack images per case in 3 cases. C. The number of cryptic splicing puncta of each target per neuron with TDP-43 pathology. The number of neurons quantified: KALRNce, n=137; RAP1GAPce, n=158; KCNQ2ce, n=236; SETD5ce, n=165; SYT7ce, n=19. D. The number of normal splicing puncta of each target per field of images from two patients and two age-matched healthy controls. (mean ± s.e.m., n=3, unpaired t-test, * p<0.05, ** p<0.01, *** p<0.001, **** p<0.0001)

**Figure S3.**
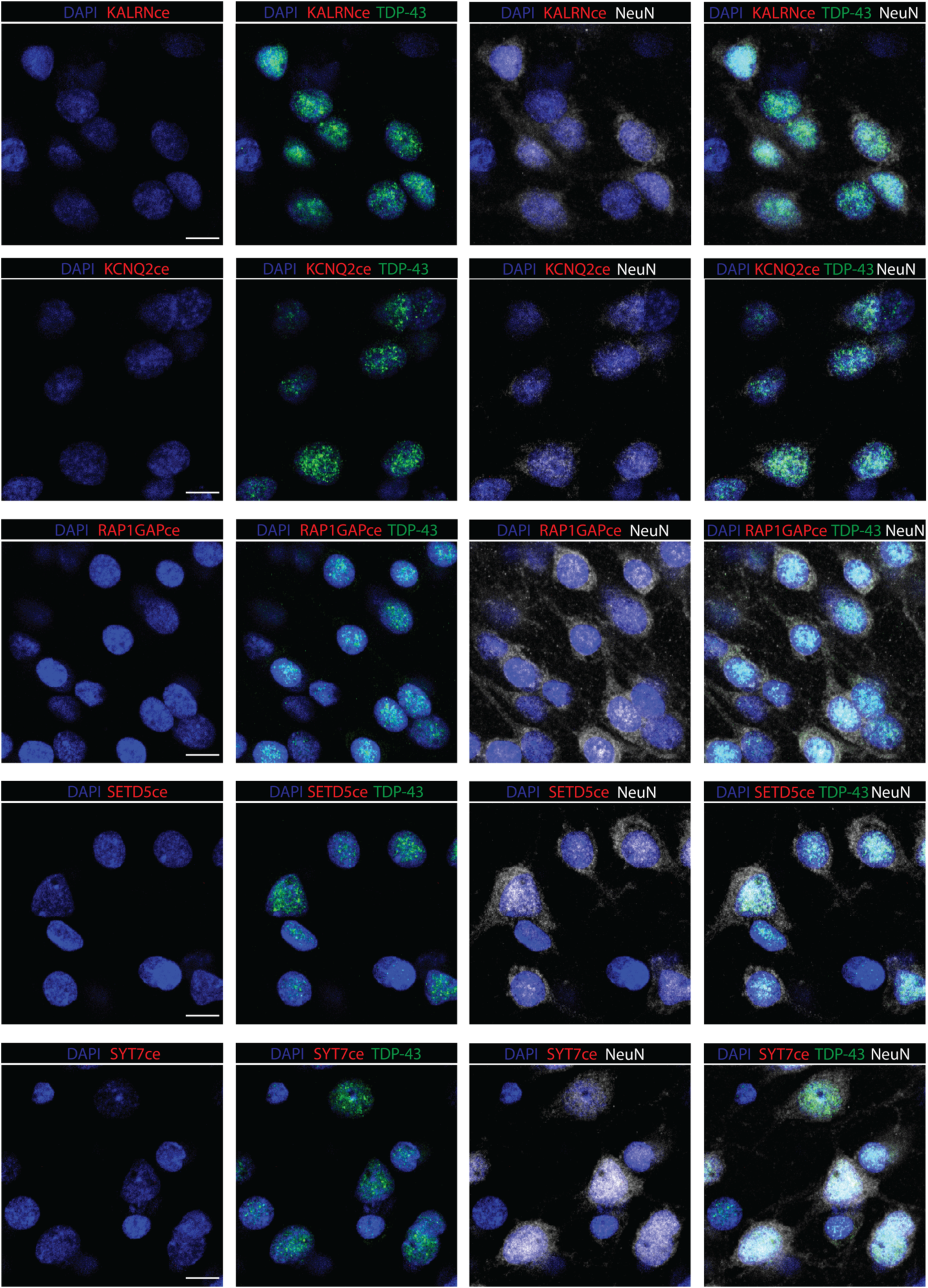
Cryptic splicing cannot be detected in healthy individuals BaseScope *in situ* hybridization shows no puncta of transcripts with a cryptic exon of *KALRN*, *KCNQ2*, *RAP1GAP*, *SETD5*, and *SYT7* (red), in combination with immunofluorescent staining of the nuclear marker DAPI (blue), TDP-43 (green), and the neuronal marker NeuN (white) on sections from the medial frontal cortex of healthy individuals who are age-matched controls to FTD-MND patients. Scale bar, 10 μm.

**Figure S4.**
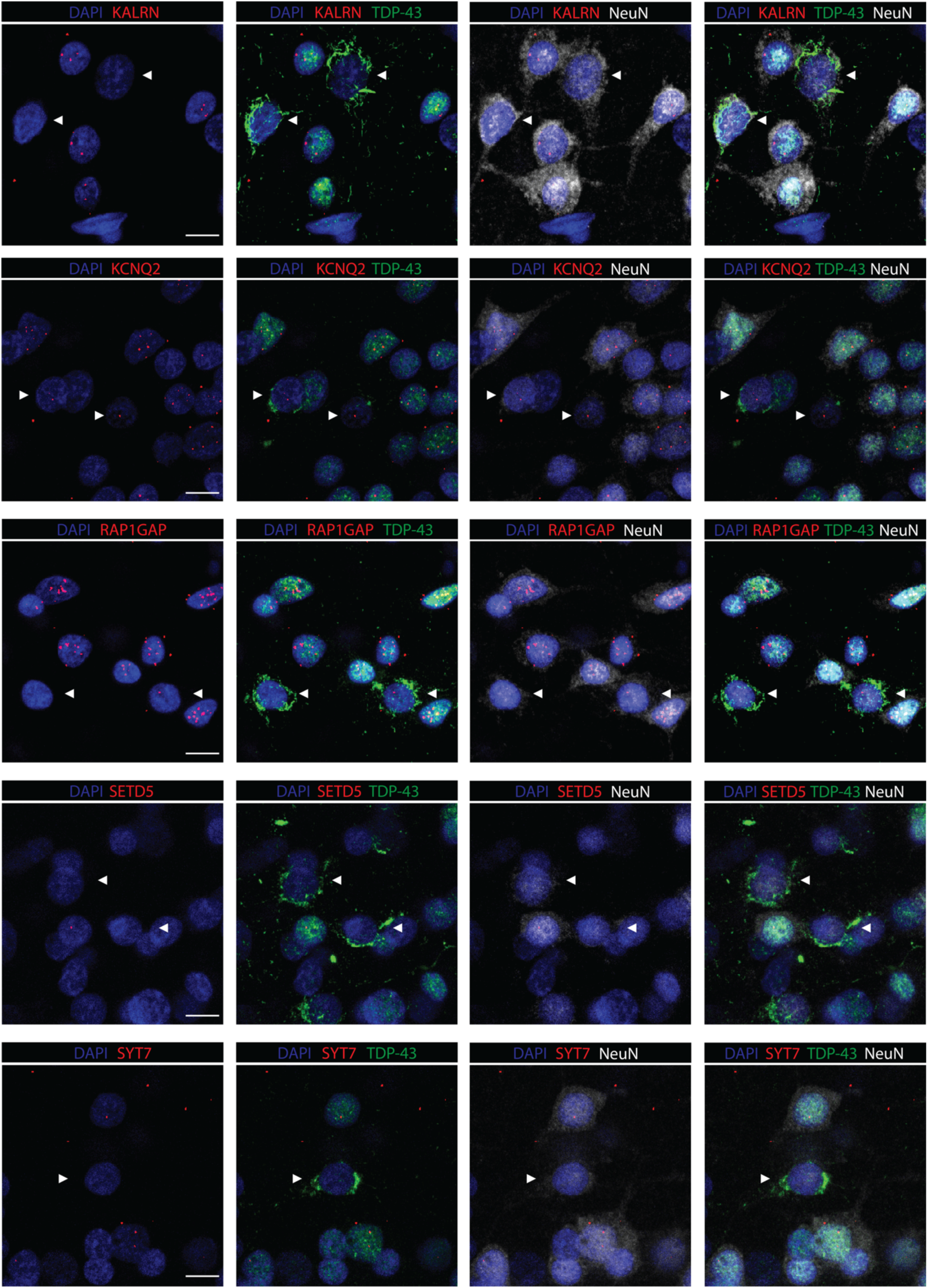
Normal splicing of each target is reduced in neurons lacking nuclear TDP-43 in brains from FTD-MND patients BaseScope *in situ* hybridization shows puncta of normal-spliced transcripts of *KALRN*, *KCNQ2*, *RAP1GAP*, *SETD5*, and *SYT7* (red), in combination with immunofluorescent staining of the nuclear marker DAPI (blue), TDP-43 (green), and the neuronal marker NeuN (white) on sections from the medial frontal cortex of FTD-MND patients. Arrowheads indicate neurons lacking nuclear TDP-43 and with cytoplasmic TDP-43 aggregates. Scale bar, 10 μm.

**Figure S5.**
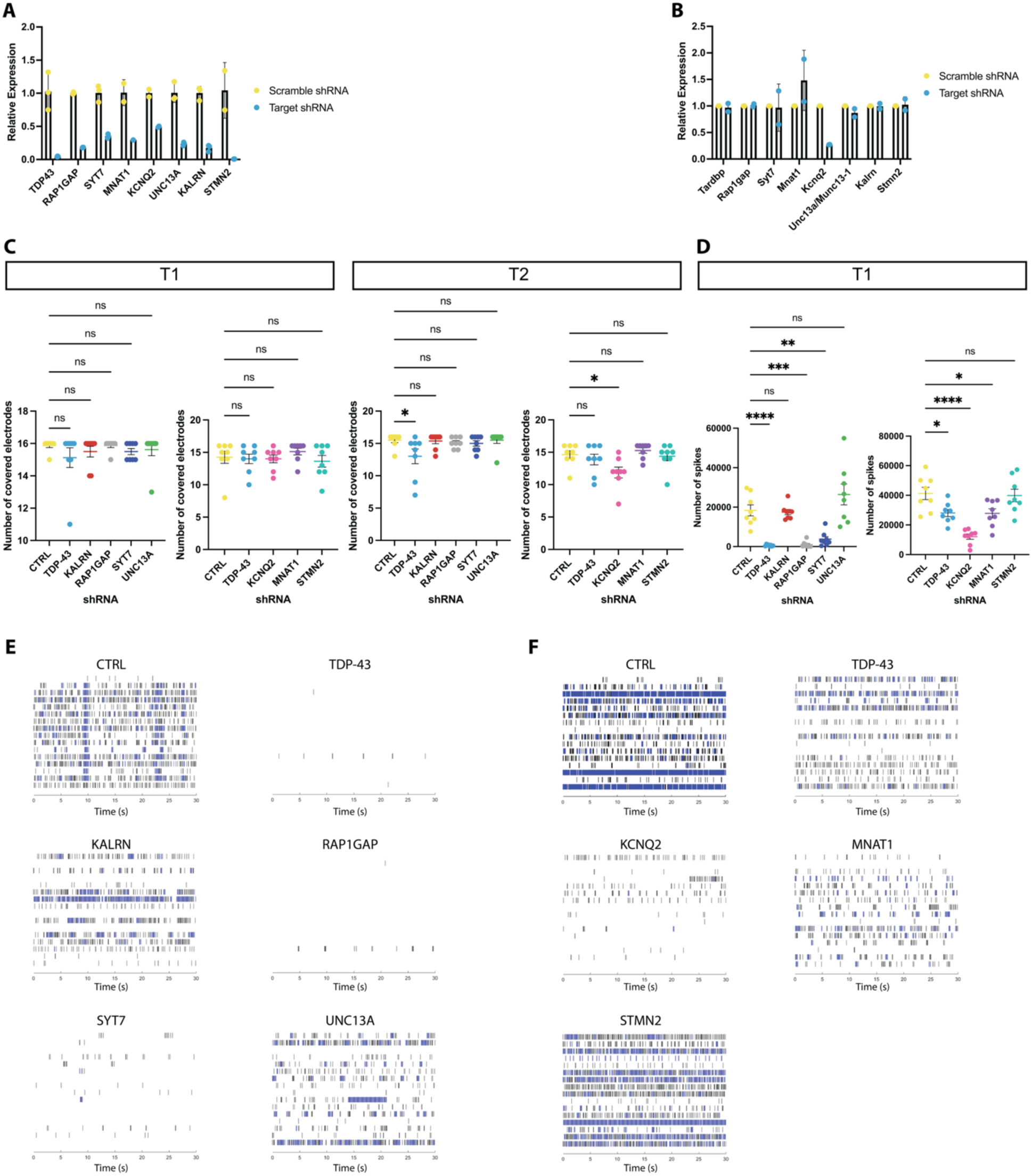
Knocking down TDP-43 or its splicing targets impairs synaptic function in iNeurons A. RT-qPCR shows decreased gene expression of TDP-43 or its splicing targets using human specific shRNAs for 7 days in iNeurons. B. RT-qPCR shows no change in gene expression of TDP-43 or its splicing targets using human specific shRNAs for 7 days in mouse primary culture, except for *KCNQ2*. C. MEA analysis reveals no decrease or mild decrease in the number of covered electrodes upon reductions in TDP-43, or its cryptic splicing targets, KALRN, RAP1GAP, SYT7, UNC13A, KCNQ2, MNAT1, and STMN2 in iNeurons. Batch 1 examined the effects of a reduction in TDP-43, KALRN, RAP1GAP, SYT7, and UNC13A, and Batch 2 examined the effects of a reduction in TDP-43, KCNQ2, MNAT1, and STMN2. Knockdown or scramble shRNAs were administered on Day 21 in both batches. A first evaluation was conducted on Day 40 in Batch 1 and on Day 30 in Batch 2 (T1). A second evaluation was conducted on Day 48 in Batch 1 and on Day 35 in Batch 2 (T2). D. MEA analysis shows the number of spikes (spontaneous firing) is decreased by the reduction in TDP-43, RAP1GAP, SYT7, KCNQ2, and MNAT1 in iNeurons at T1. E. Representative raster plots show neuronal activity in a 30-second time frame for each knockdown condition in Batch 1. F. Representative raster plots show neuronal activity in a 30-second time frame for each knockdown condition in Batch 2.

**Figure S6.**
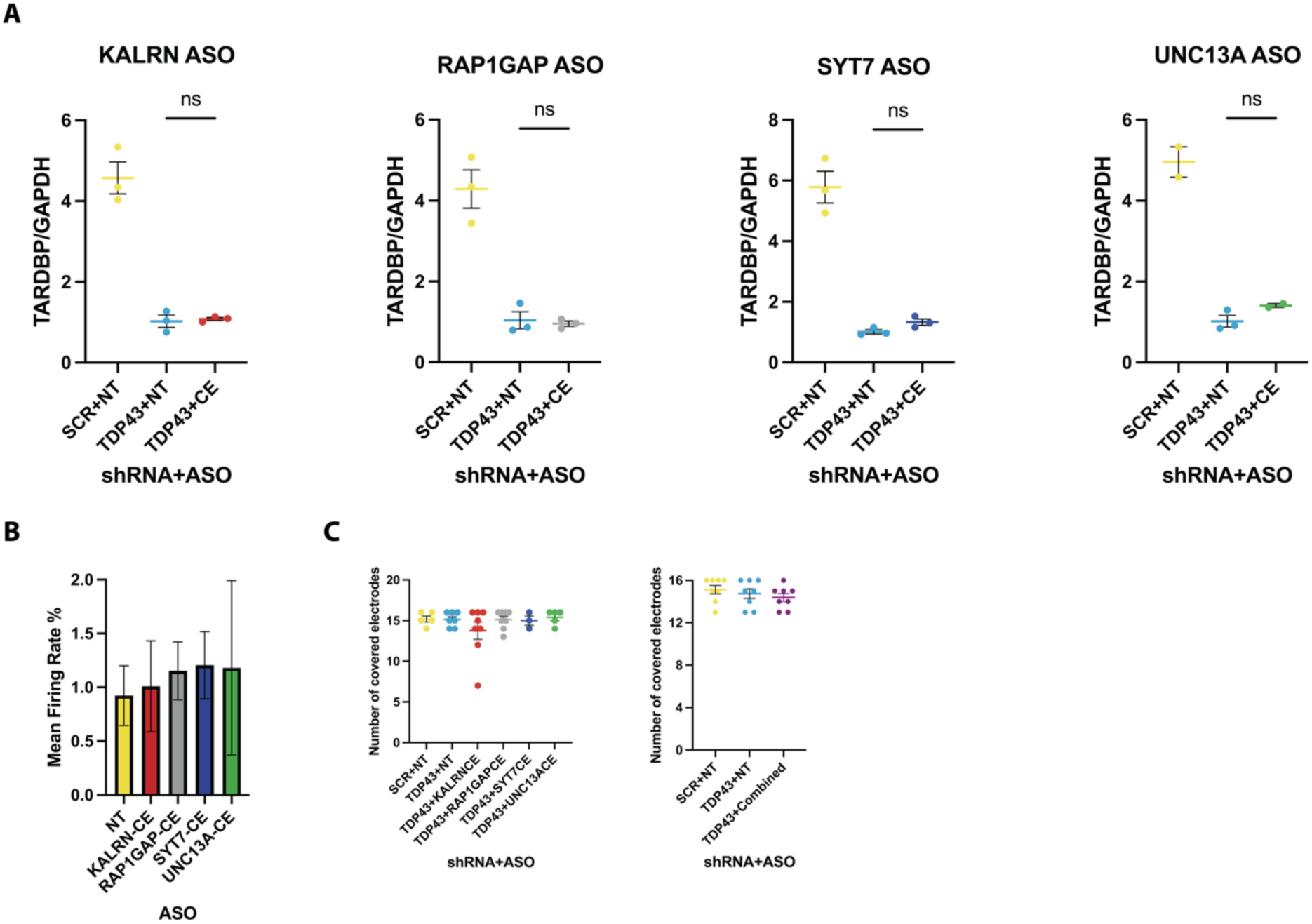
Splicing-inhibiting ASO treatment does not change TDP-43 level upon TDP-43 reduction or neuronal activity under normal condition A. RT-qPCR shows decreased gene expression of TDP-43 when iNeurons treated with TDP-43 shRNAs, which does not change upon ASO treatment. B. MEA analysis reveals no difference in the normalized mean firing rate upon ASO treatment under normal condition. The normalized mean firing rate is calculated dividing the mean firing rate on Day 35 by the mean firing rate on Day 20 when ASO treatment started. C. MEA analysis reveals no decrease in the number of covered electrodes upon reductions in TDP-43 and ASO treatment.

**Supplementary Table 1:**
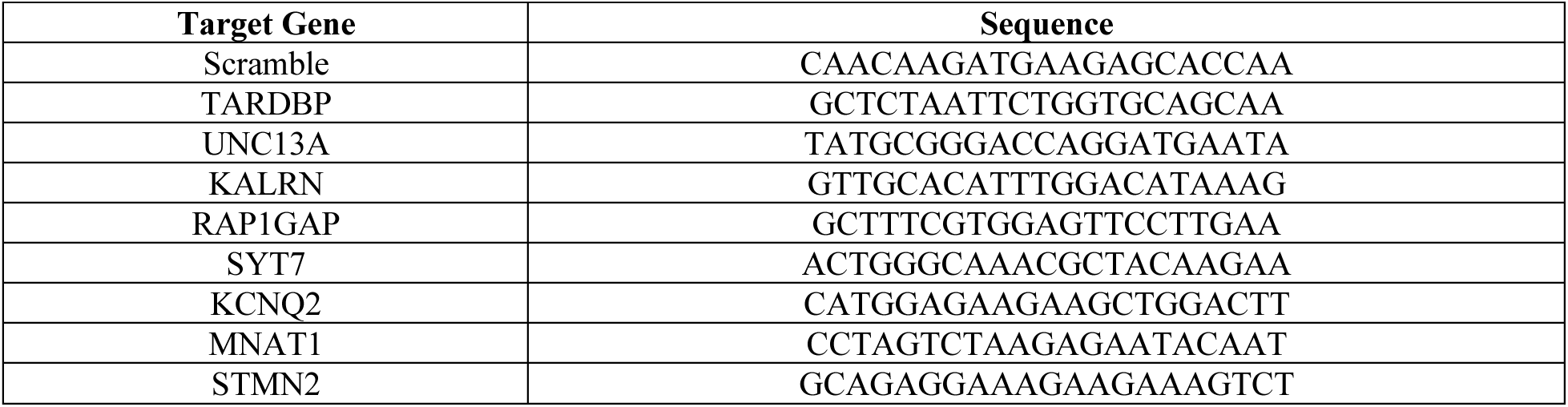
List of shRNA sequences used in this study.

**Supplementary Table 2:**
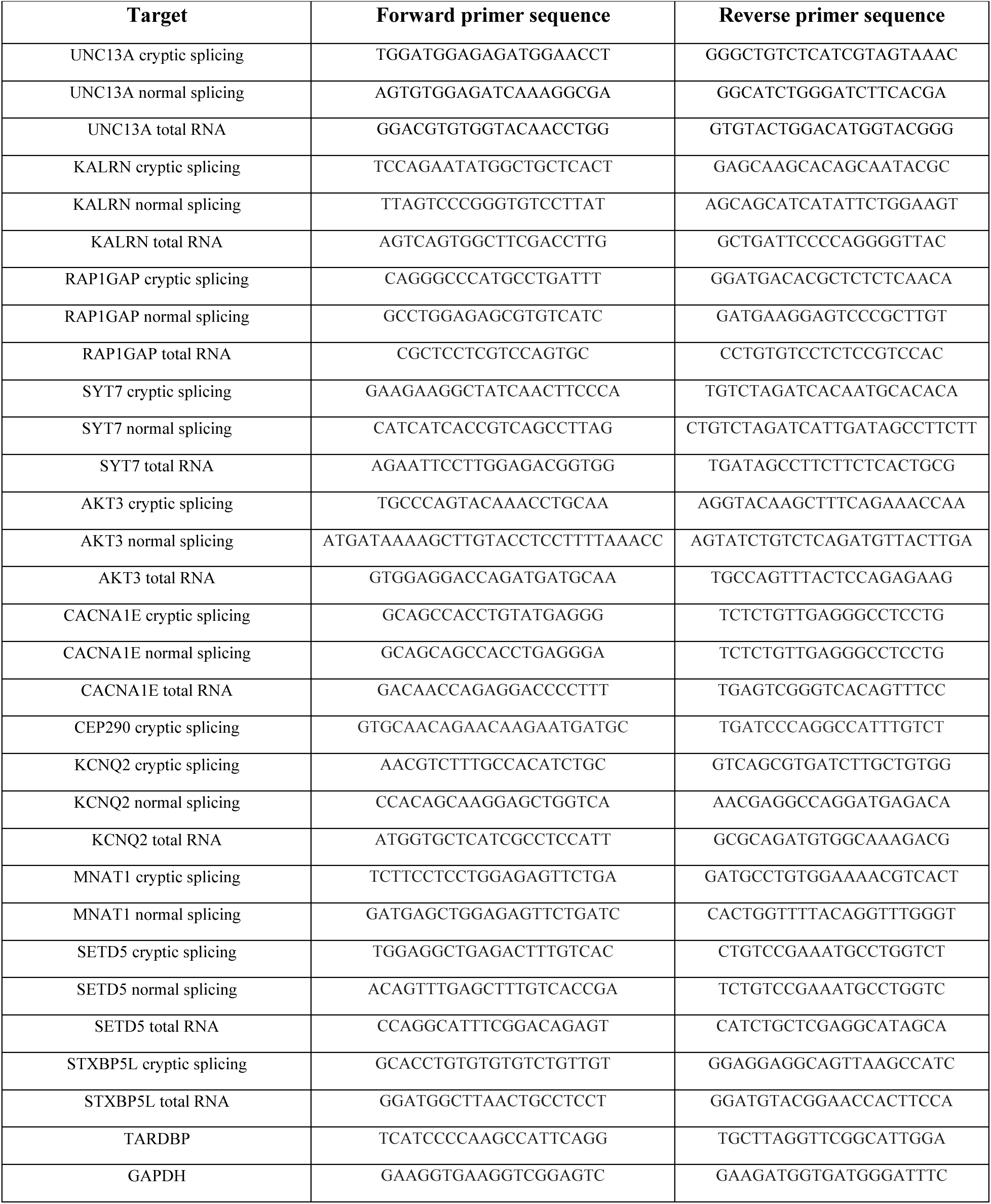
List of primer sequences used in this study.

**Supplementary Table 3:** List of sporadic FTLD-TDP and control cases used for the RNA *in situ* hybridization experiment (ISH) and the RNA extraction and qPCR experiment (qPCR)

| Case No. | Age (years) | Sex | PMI (hrs) | Clinical diagnosis | Primary neuropathological diagnosis | ADNC | ISH or qPCR |
| --- | --- | --- | --- | --- | --- | --- | --- |
| FTD-MND 1 | 66 | M | 12.1 | bvFTD-MND | FTLD-TDP-B, ALS | Low | ISH |
| FTD-MND 2 | 72 | M | 6.7 | bvFTD-MND | FTLD-TDP-B | Not | ISH and qPCR |
| FTD-MND 3 | 57 | M | 7.6 | bvFTD-ALS | FTLD-TDP-B, MND | Not | ISH and qPCR |
| FTD-MND 4 | 65 | F | 8.5 | bvFTD | FTLD-TDP-B, MND | Low | qPCR |
| Control 1 | 76 | M | 8.2 | Control | None | Low | ISH and qPCR |
| Control 2 | 60 | F | 20.5 | Control | None | Low | ISH |
| Control 3 | 72 | M | <19 | Control | None | Low | ISH |
| Control 4 | 86 | F | 7.8 | Control | None | Low | qPCR |
| Control 5 | 78 | F | 11.6 | Control | None | Low | qPCR |

Abbreviations: PMI – postmortem interval, bvFTD – behavioral variant frontotemporal dementia, ALS – amyotrophic lateral sclerosis, MND – motor neuron disease, FTLD – frontotemporal lobar degeneration, ADNC - Alzheimer’s disease neuropathologic change

**Supplementary Table 4:**
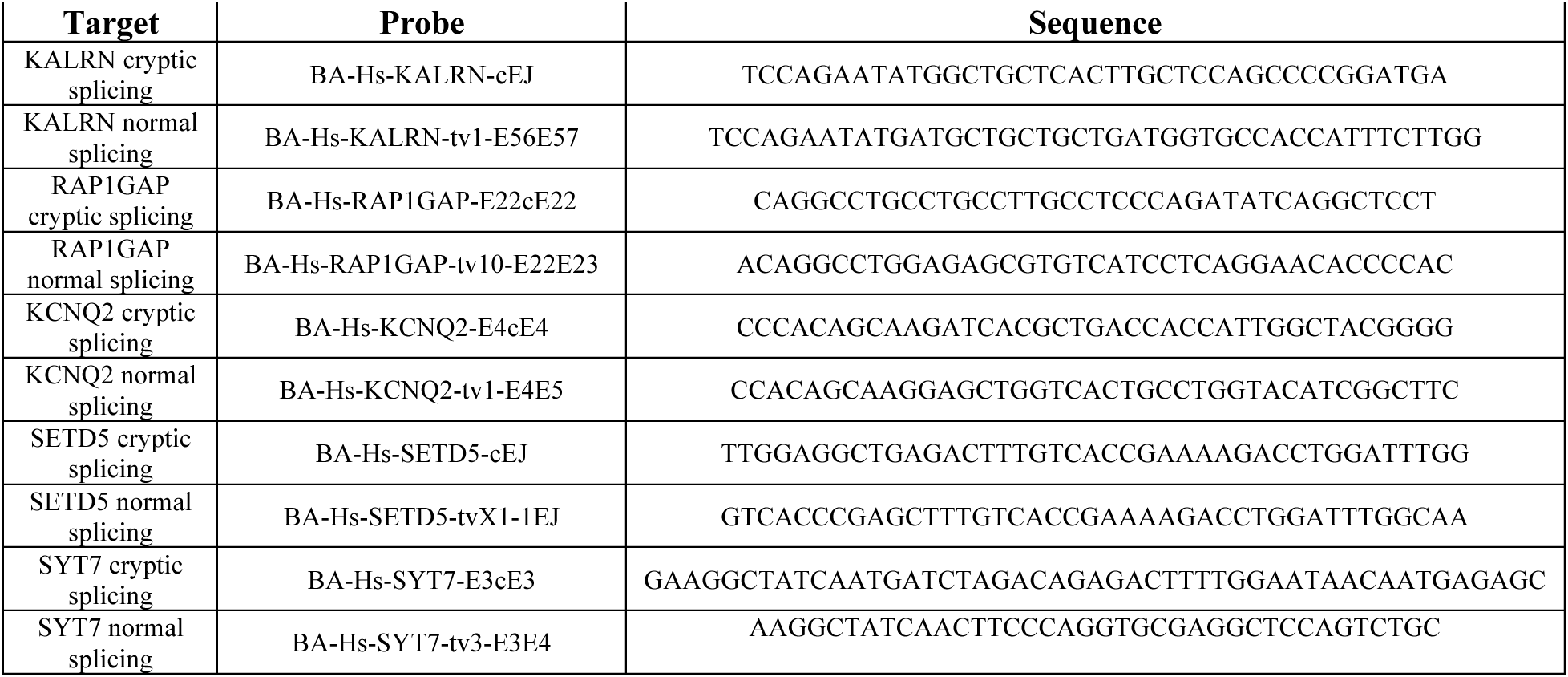
List of probe sequences for the RNA *in situ* hybridization experiment.

